# Assessing and assuring interoperability of a genomics file format

**DOI:** 10.1101/2022.01.07.475366

**Authors:** Yi Nian Niu, Eric G. Roberts, Danielle Denisko, Michael M. Hoffman

## Abstract

**Background:** Bioinformatics software tools operate largely through the use of specialized genomics file formats. Often these formats lack formal specification, and only rarely do the creators of these tools robustly test them for correct handling of input and output. This causes problems in interoperability between different tools that, at best, wastes time and frustrates users. At worst, interoperability issues could lead to undetected errors in scientific results.

**Methods:** We sought (1) to assess the interoperability of a wide range of bioinformatics software using a shared genomics file format and (2) to provide a simple, reproducible method for enhancing inter-operability. As a focus, we selected the popular Browser Extensible Data (BED) file format for genomic interval data. Based on the file format’s original documentation, we created a formal specification. We developed a new verification system, Acidbio (https://github.com/hoffmangroup/acidbio), which tests for correct behavior in bioinformatics software packages. We crafted tests to unify correct behavior when tools encounter various edge cases—potentially unexpected inputs that exemplify the limits of the format. To analyze the performance of existing software, we tested the input validation of 80 Bioconda packages that parsed the BED format. We also used a fuzzing approach to automatically perform additional testing.

**Results:** Of 80 software packages examined, 75 achieved less than 70% correctness on our test suite. We categorized multiple root causes for the poor performance of different types of software. Fuzzing detected other errors that the manually designed test suite could not. We also created a badge system that developers can use to indicate more precisely which BED variants their software accepts and to advertise the software’s performance on the test suite.

**Discussion:** Acidbio makes it easy to assess interoperability of software using the BED format, and therefore to identify areas for improvement in individual software packages. Applying our approach to other file formats would increase the reliability of bioinformatics software and data.

## 1 Introduction

### 1.1 File format interoperability

For your latest research project, you have constructed a pipeline from multiple published bioinformatics tools. Each tool works well with the author’s data, but you run into errors with your data. The author’s data and your data have slight differences in file metadata and data formatting, which lead to the errors. As a result, you must spend time manually editing your data files and intermediate outputs to conform to each tool’s expectations. Meanwhile, ensuring interoperability between software tools that parse the data file format could have prevented your frustration.

Scientific software developed by academics often suffers from software engineering deficiencies^1^, which can lead to the scenario described above. Among these include problems with deployment^2^, maintenance^3^, robustness^4^, and documentation^5^. Software engineering flaws may hinder fulfilling the Findable, Accessible, Interoperable, and Reusable (FAIR) principles for scientific data management^6^—especially the guidelines on interoperability and reusability. Software engineering flaws may also affect web services that parse bioinformatics file formats, which may have vulnerabilities to attacks such as malicious code injections in input files^7^.

One key difficulty arises from interoperability of specialized file formats used for scientific data. Often, creators specify such formats informally, or not at all, leaving users and developers to guess the details of critical components or edge cases. Rare standardization efforts such as those of the Global Alliance for Genomics and Health (GA4GH)^8^ have developed a few formal specifications. These include the sequence alignment/map (SAM), BAM, CRAM, and variant call format (VCF) file formats^9^.

Interoperability issues can also arise from issues within the software. Developers can address some interoperability problems, however, through simple solutions such as checklists. For example, Bioconda^10^ recipes require adequate tests and a stable source code uniform resource locator (URL)^11^. Bioconductor^12^ also has guidelines for package submission regarding code style, performance and testing^13^. Simple checklists can greatly improve software quality, even for programmers and researchers that lack formal software engineering training.

Software testing recommendations and standard test suites can aid researchers and developers. Extensive test suites for common standards, such as TeX’s trip tests^14^, or the Web Standards Project Acid test suite^15^ exercise independent implementations of common standards by focusing on edge cases. In a bioinformatics context, tools that parse the VCF format^16^ can use simulated VCF files with known behavior to test software correctness^17^.

Here, we tackled the bioinformatics software engineering problem of file format interoperability, specifically focusing on the plain-text whitespace-delimited Browser Extensible Data (BED) format^18^. We chose to use the BED file format because of its simplicity and its popularity. First, we developed a formal specification for the BED format as a comprehensive specification did not exist. Second, we quantified the degree to which a wide variety of bioinformatics software varied in their processing of this file format. In particular, we tested bioinformatics software input validation, checking input data for correct formatting. To facilitate this work, we created Acidbio (https://github.com/hoffmangroup/acidbio), a system for automated testing and certification of bioinformatics file format interoperability.

### 1.2 The BED file format

The BED format describes genomic intervals in plain text. Each BED file consists of a number of lines, each with 3 to 12 whitespace-delimited fields. The mandatory first three fields (chrom, chromStart, and chromEnd) define an interval on a chromosome. The optional last nine fields provide additional information about the interval such as a name, score, strand, and aesthetic features used by the University of California, Santa Cruz (UCSC) Genome Browser^19^. The optional fields have binding order—all fields preceding the last field used must contain values.

BED variants distinguish BED files based on its number of fields. BED*n* denotes a file with only the first *n* fields. For example, a BED4 file has the chrom, chromStart, chromEnd, and name fields. BED3 to BED9, along with BED12, represent the 8 standard BED variants.

BED*n*+*m* denotes a file with the first *n* fields followed by *m* fields of custom-defined fields supplied by the user. The custom-defined fields can contain many types of plain-text data. BED*n*+*m* files act as custom BED files. Currently, no in-band information exists to supply information about a BED file’s fields. A BED parser must infer the fields present in a BED file.

The file conversion tool bedToBigBed^18^, developed by the UCSC Genome Browser team^20^, has served as the de facto file validation tool for the BED format. The BED format appears deceptively simple, and without careful consideration of the specification, a developer may miss unexpected flexibility or rigidity in some fields.

## 2 Results

### 2.1 A new formal specification addresses ambiguities in the BED format

Despite existing for almost two decades, the BED format until recently lacked a formal specification similar to the SAM^21^ or VCF^16^ specification. The UCSC Genome Browser Data File Formats Frequently Asked Questions (https://genome.ucsc.edu/FAQ/FAQformat.html) specified some details, but lacked technical details that other formal specifications clearly define.

Through the GA4GH standards process^8^, we established a specification of the BED format (https://github.com/samtools/hts-specs/blob/master/BEDv1.pdf). The new specification defines each BED field and their possible numerical range or valid character patterns. It also provides semantics surrounding whitespace, sorting, and default field values. The specification formalizes missing details and captures the existing use of the BED format, taking the input from relevant stakeholders into account. During the development of the specification, we solicited input from a number of stakeholders, including the UCSC Genome Browser team, the File Formats subgroup within the GA4GH Large Scale Genomics work stream (https://www.ga4gh.org/work_stream/large-scale-genomics/), and the public through GitHub comments (https://github.com/samtools/hts-specs/pull/570).

### 2.2 Most existing tools perform poorly on a BED test suite

To measure the ability of BED parsers to accept good input and reject bad input, we created an Acidbio test suite with 92 individual test cases. Specifically, we used the new specification to develop a test suite of expected pass and expected fail BED files. The expected pass test cases conform to our specification—for these cases, we expect tools to return a zero exit code and not output any error or warning messages. The expected fail test cases do not conform to our specification—for these cases, we expect tools to return a non-zero exit code or output an error or warning message. The test suite contains 92 tests, covering the definitions of fields and the structure of the BED file. The test suite also covers all BED variants from BED3 to BED12. The BED3 test cases represent the core of our test suite, as all BED files must have the first three fields.

The BED format does not contain in-band information on whether a file uses BED fields only or also has custom fields. A parser might assume that for BED files with 4 to 12 fields, all the fields represent standard BED fields. In this case, the parser should validate the fields according to the file format rules.

Alternatively, a parser might treat fields 4 through 12 as custom data. A tool designed to handle arbitrary custom BED files may not validate the optional BED fields. This means the tool may not fail on the expected fail test cases. The expected success test cases, however, should all work even for non-specified custom data. Also, this flexibility does not apply to mandatory fields 1 through 3, as their definition cannot change.

We examined behavior of tools, expecting strict validation of standard BED4 through BED12 files. This provides more informative results than permitting the whole range of behavior one might expect for custom data. Unexpected results in the optional fields indicate the need for better means for interchange of metadata on these fields.

Using our test suite, we assessed 80 Bioconda packages that support the BED format as input (Figure 1). In some packages, we assessed multiple tools, making 99 tools in total. For each tool, we calculated its performance on each BED variant by taking the number of tests that behaved as expected divided by the number of tests for the BED variant. Of the 99 tools, only 26 achieved ≥ 70% expected results for BED3 tests. Averaged for tests across all BED variants, 51 tools achieved ≥ 50% expected results. We have deposited full results on Zenodo (https://doi.org/10.5281/zenodo.5784787). Beyond the possibility of expecting custom BED files, we attributed unexpected results to several causes described below.

**Figure 1:**
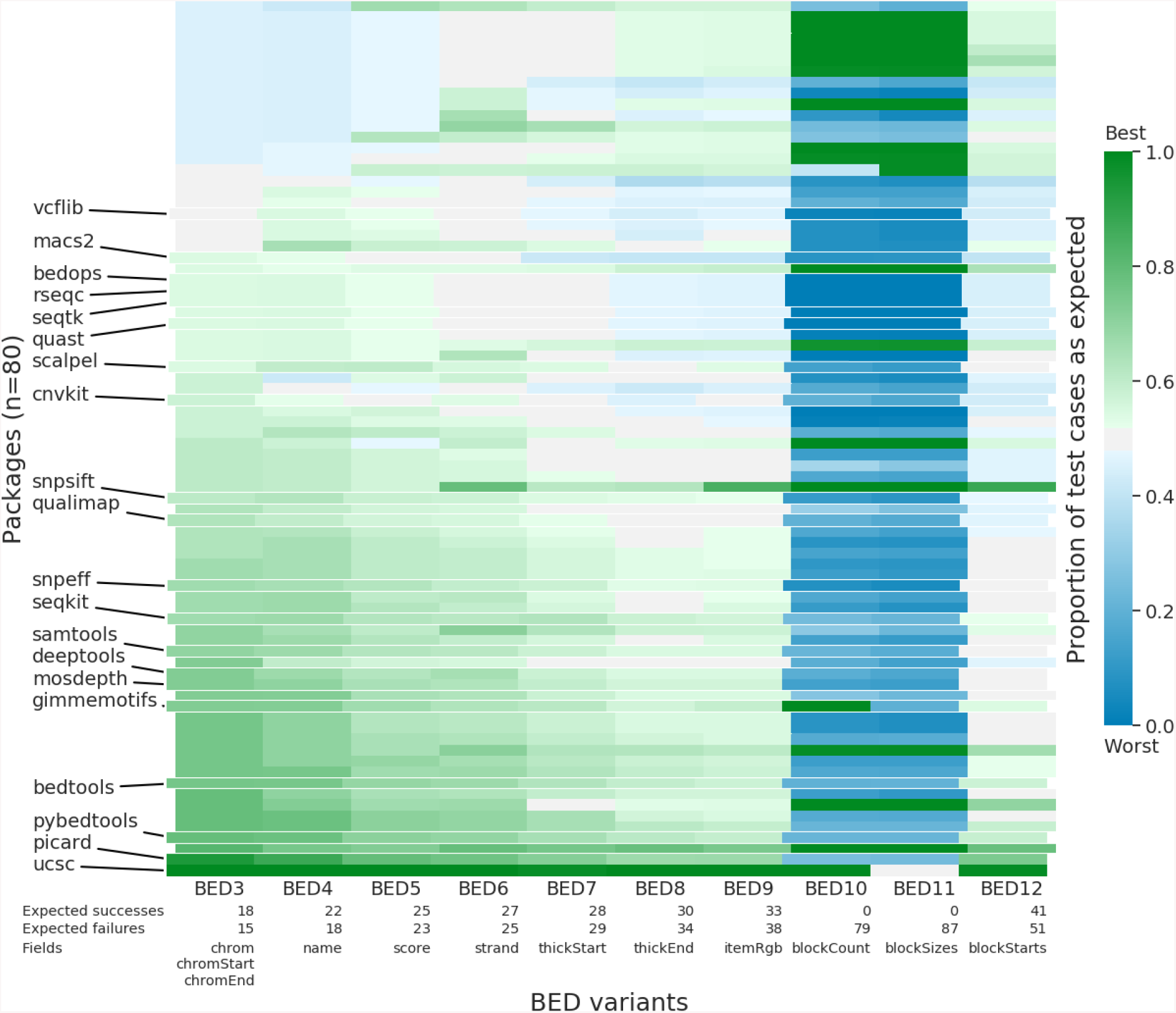
Heatmap of performance of 80 Bioconda packages on 10 BED variants. Each cell shows the percentage of successful tests from the BED variant. Green cells: strong performance on the test suite; blue cells: poor performance. An expected success test case that succeeds or an expected fail test case that fails both represent a successful test. For packages with multiple tools, we display only results from the package’s best-performing tool. Labeled, negative indented rows emphasize the 20 packages most downloaded from Bioconda. Rows sorted by ascending performance on the mandatory BED3 fields, then on performance of subsequent optional fields, ending with BED12 performance. The table below the heatmap lists the number of expected success and expected fail test cases for each BED variant. BED10 and BED11 have zero expected success test cases because the specification forbids BED10 and BED11.

### 2.3 Existing tools parse BED files in different ways

All tools have distinct purposes, causing them to parse the BED format in different ways and focus on varying aspects of BED files. Different purposes mean some test cases may never arise in the expected usage of the tool. We have identified a few groups of tools that have similar behaviors, which cause poor performance on the test suite.

#### Tools that require a specific BED variant

Some tools require a specific number of fields in the input BED file. For example, slncky^22^ requires a BED12 file. This causes all BED3 to BED11 inputs to raise an error.

#### Tools that only validate a subset of BED fields

Many tools use the BED format only for interchange of genomic intervals in the first three fields. Some of these tools will accept any BED *n* file and perform no validation after the first three fields. For example, many tools ignore fields that describe aesthetic features only for genomic browser display, such as thickStart, thickEnd, and itemRgb. A tool such as bedtools^23^ that mainly operates on genomic intervals would incorrectly succeed on an expected fail BED9 test case.

#### File converters

Some tools convert the BED format to a different file format, without performing any validation. Some file converters use a garbage-in-garbage-out approach, going from invalid input in BED format to invalid output in some other format. For example, bioconvert bed2wiggle^24^ fails as expected on most expected fail test cases, but still produces output retaining the input file errors. Using a garbage-in-garbage-out approach may make debugging complex pipelines more difficult. Raising warnings during file conversion helps debugging, as the user can narrow down the source of the error to steps before file conversion.

#### Tools that use another library for BED parsing

Some tools call an external library to perform operations on BED files. If the main tool does not perform extra error checking of its own, it can only detect the same errors that the external library finds. For example, intervene^25^ uses bedtools as a dependency, which results in their similar patterns of performance.

### 2.4 Ambiguous format specification makes uniform behavior more difficult

The previous absence of a formal specification for the BED format also influenced test performance. Our formal specification and the behavior of the reference implementation bedToBigBed conflict with the expectations of tool developers in many ways.

#### Definition of whitespace

Many BED files use tabs to delimit fields. The BED format, however, also accepts spaces to delimit fields, if the fields themselves contain no spaces^20^. Of the 99 tools examined, 60 reject space-delimited BED files allowed by the specification (Table 1, “other-fully_space_delimited.bed”). Also, the BED format permits blank lines, though 37 tools do not accept this (Table 1, “other-space_between_lines.bed”).

**Table 1:** Number of tools that passed each of 18 expected success BED3 test cases. We tested 99 tools. The leading number in the test case name describes the BED field that the test focuses on. “Other” means the test focuses on the structure of the file, rather than a particular field.

| Tools passing | Test case |
| --- | --- |
| 37 | other-fully_space_delimited.bed |
| 37 | other-not_tab_delimited.bed |
| 52 | other-comment_start.bed |
| 57 | other-space_before_tab.bed |
| 58 | other-space_after_tab.bed |
| 58 | other-space_between_lines.bed |
| 60 | other-multiple_comments.bed |
| 61 | other-comment_end.bed |
| 62 | other-comment_middle.bed |
| 67 | 03-start_eq_end.bed |
| 71 | 01-known_scaffolds.bed |
| 71 | 01-unknown_scaffolds.bed |
| 78 | 02-leading_zeros_start.bed |
| 79 | 03-duplicate_rows.bed |
| 79 | 03-overlapping_intervals.bed |
| 79 | 03-zero_chromStart.bed |
| 79 | 03-no_gap_startend.bed |
| 79 | 03-standard_chrom.bed |

#### Expanded definition of fields

The BED format requires strict limits for certain fields and some generators do not respect these limits. For example, the specification defines score as an integer value between 0 and 1000, inclusive. Some tools use the score as a p-value, which violates the integer definition. To allow tools to repurpose the nine optional fields, one can treat these tools as BED*n*+*m* parsers, with custom definitions for the remaining fields. Nonetheless, repurposing field names, such as score, with different definitions can confuse parsers that will misinterpret the data and use it incorrectly.

#### Conflict between our formal specification and bedToBigBed

We used the de facto file validator bedToBigBed to inform the design of our test suite. Without a formal specification, however, uncertainty surrounding specific edge cases arose when bedToBigBed disagreed with our understanding of correct behavior.

Our formal specification disagreed with bedToBigBed in three instances. First, bedToBigBed accepted a BED7 file with thickStart less than chromStart. Second, bedToBigBed accepted a BED12 file with the length of the blockSizes or blockStarts list greater than blockCount. Third, bedToBigBed accepted BED11 files while our specification disallowed BED11.

### 2.5 Software engineering deficiencies lead to poor performance on the test suite

Beyond issues in differences in design between tools and the previous informal specification of the file format, we can also attribute poor testing performance to problems in software engineering.

#### Silently accepting invalid input

Tools should alert users on input errors, allowing them to check whether they have made an error. In some cases, developers prefer to skip an invalid data point and continue. In this case, the tool should at least provide a warning message describing the skipped line. Otherwise, an error could slip past the user and affect their results. In our test suite, a warning message would count as an expected failure, improving the performance statistics for a tool that generates them.

Errors in BED file generators can easily slip past users. When a downstream tool raises an error on bad input, this reduces the time before someone discovers the problem with the upstream generator.

#### Insufficient testing

While some of our test cases cover formatting issues that can hinder interoperability, others represent “can’t happen” scenarios that, uncaught, pose logic bombs for a software tool. For example, all tools should reject negative start positions (Figure 2), “02-negative-start.bed”, but 48/99 tools accepted a test case that has negative starts. Given the limited resources and incentives to publish in academic software engineering, developers require a simpler way to ensure avoidance of obvious problems than manually developing test cases.

**Figure 2:**
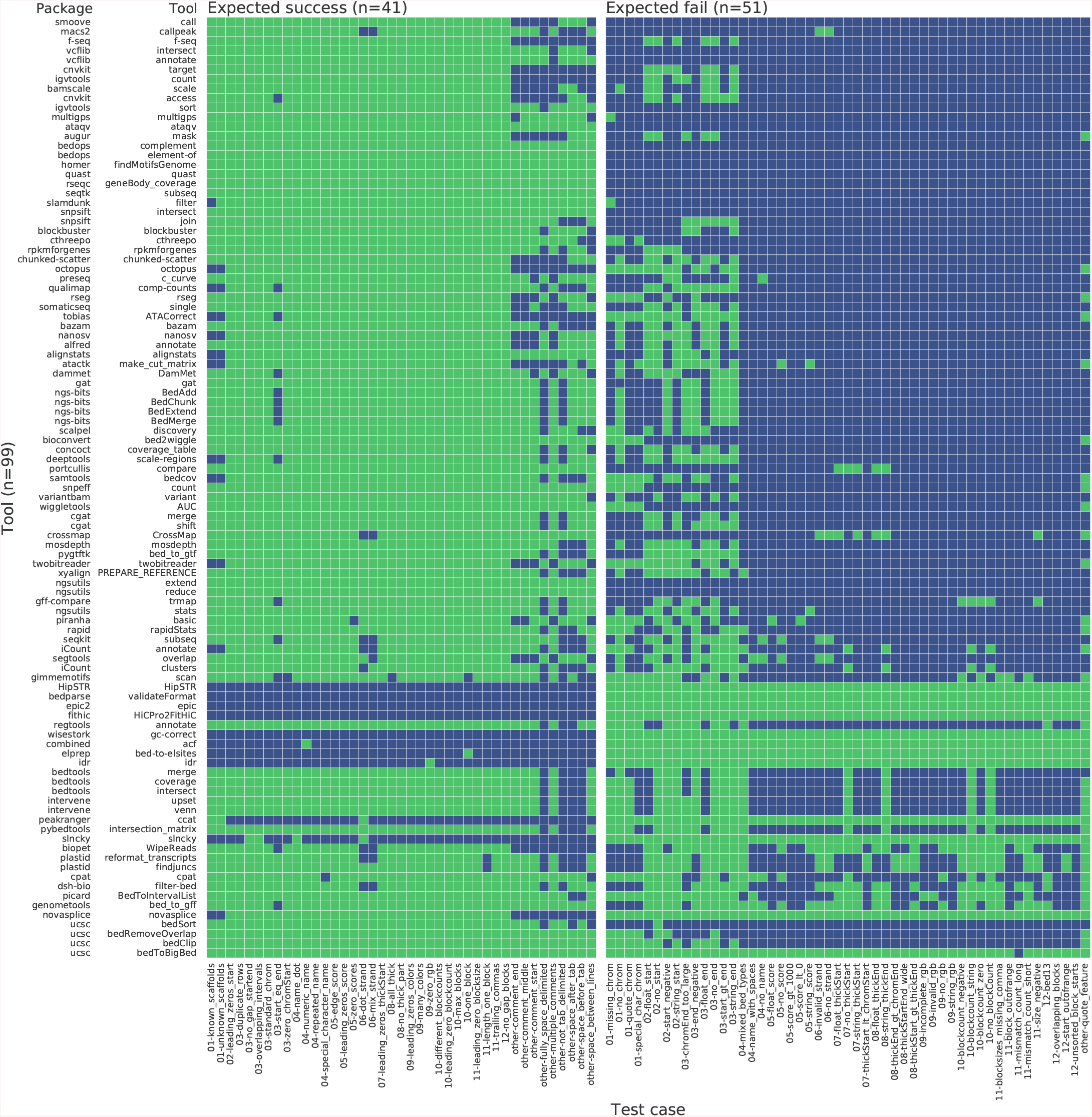
Performance of 99 tools from 80 packages on 92 BED12 test cases^21–106^. Green: the tool performed as expected; blue: the tool did not perform as expected. Rows sorted ascending by the number of test cases with an expected result. We grouped tools in the same package together as they tend to have similar results. For packages with multiple tools, we sorted the package using the best-performing tool. Within the same package, we sorted tools by ascending performance.

### 2.6 No relationship between package performance and downloads found

We observe little correlation between the number of downloads a package has compared to the package’s performance on the test suite (Figure 3). Many packages have a similar number of downloads. We attribute this to packages having specific purposes that make them useful for a few users. However, very highly downloaded packages such as bedtools^23^ and the UCSC Genome Browser tool suite^83^ have better performance than other tools.

**Figure 3:**
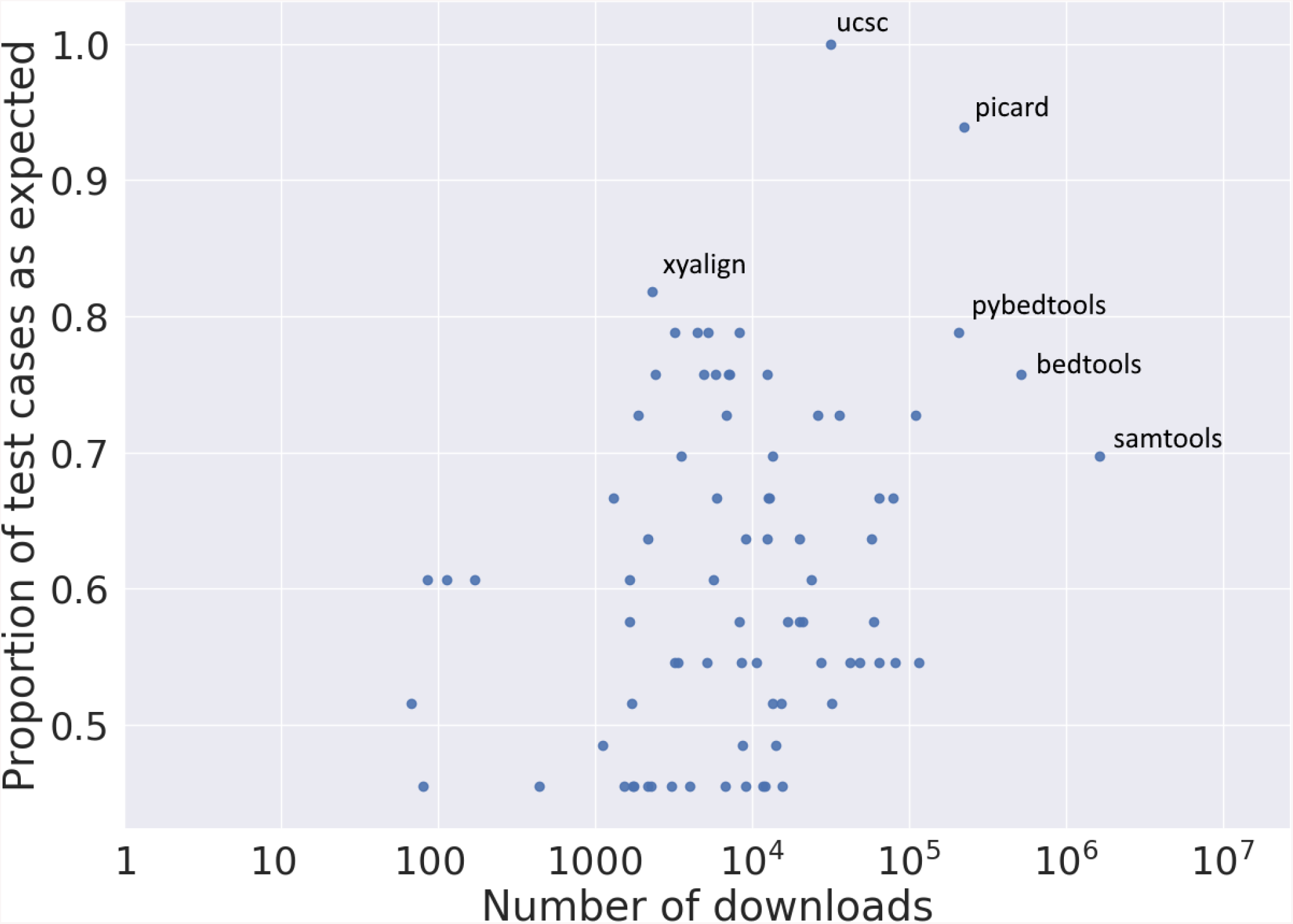
Scatter plot of the number of downloads a package has on Bioconda against its performance on BED3 tests. Labeled points indicate the top 4 performing tools and the top 4 most downloaded tools. For packages with multiple tools, we display results from the best-performing tool.

### 2.7 Automated fuzzing can detect errors that a manually designed test suite does not

Differential testing^107^ using files generated from a grammar-based fuzzer^108^ can discover new errors not found by the test suite. A grammar-based fuzzer automatically generates files based on a defined structure of the file format.

We found one example of unexpected behavior in bedtools coverage^23^ where coverage raised an error but bedToBigBed did not. Since bedtools coverage requires two input files, we generated two files using the fuzzer (Table 2) and validated them using bedToBigBed. On the generated files, bedtools coverage exited with exit status 1 and error message “Error: line number 1 of file 2.bed has 4 fields, but 0 were expected.” Our manually designed test suite did not catch this error—we only uncovered it due to the use of fuzzing.

**Table 2:** Two files generated by a grammar-based fuzzer.

| File 1 | File 2 |
| --- | --- |
| chr18 455914 533415 woG | chr12 632184 753365 Vx6 |
|  | #I |
|  | #_ |
|  | #_ |

### 2.8 BED badge indicates conformance with the BED format

We designed badges that developers can display in a tool’s documentation to clearly indicate the file types used and indicate the tool’s performance on the test suite (Figure 4). The badges reassure users that the software underwent thorough testing. The availability of such badges encourages developers to perform input validation.

**Figure 4:**
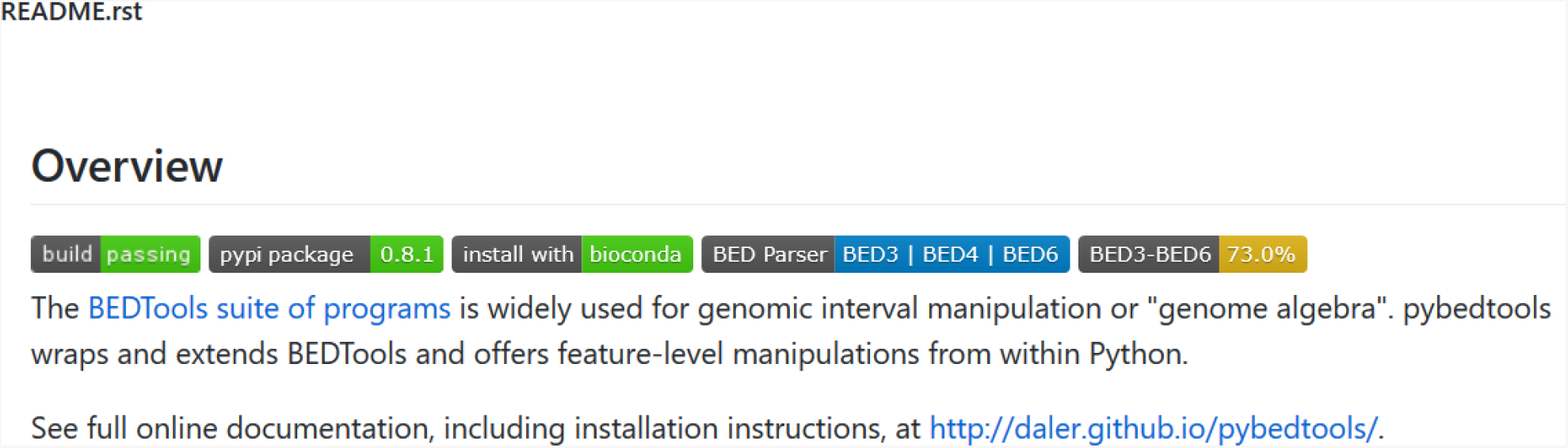
Example of BED badges. BED badges allow developers to indicate the tool’s support for the BED format and its conformance to the specification. The fourth badge, “BED parser”, displays the BED*n* formats the tool supports. The fifth badge, “BED3-BED6”, displays the average performance of the tool. In this example, the badge displays the performance averaged over BED3 to BED6, inclusive.

Acidbio includes steps to produce a BED badge. We recommend developers to display a BED badge if their software conforms to the BED formal specification.

## 3 Methods

### 3.1 The Acidbio test system

We developed the Acidbio test system, which automatically runs a number of bioinformatics tools on a test suite (Figure 5). To determine an actual success or failure, we consider the exit status and outputs to standard output and error. A test case passes on a successful exit status and no error or warning messages printed.

**Figure 5:**
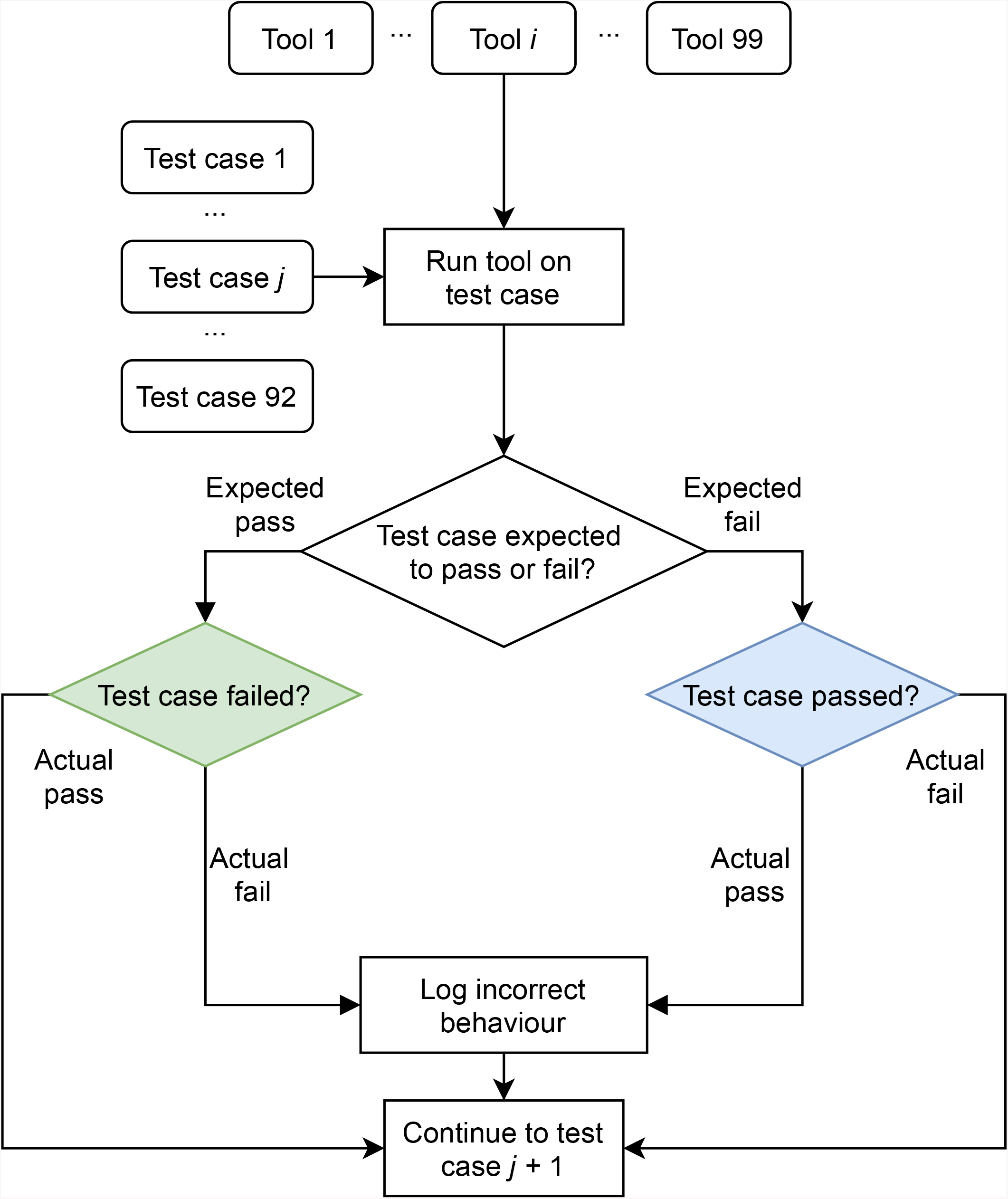
Flowchart depicting how Acidbio evaluates tools on the test suite. Rounded rectangles: inputs; sharp rectangles: procedures; rhombuses: conditional branches. All 99 tools run on all 92 test cases. After the 92nd test case, Acidbio moves onto the next tool and runs it on the first test case.

We identified error and warning messages by manually running the tools. We had to identify these error and warning messages manually because some tools logged errors without returning a non-zero exit code or logged issues in the BED file through warnings instead of errors.

To provide Acidbio with details on how to run each tool, we created a YAML Ain’t Markup Language (YAML) configuration file that stored each tool’s command-line usage file (Figure 6). The YAML file also stored the locations of the additional files needed to run each tool and each tool’s Conda environment.

**Figure 6:**
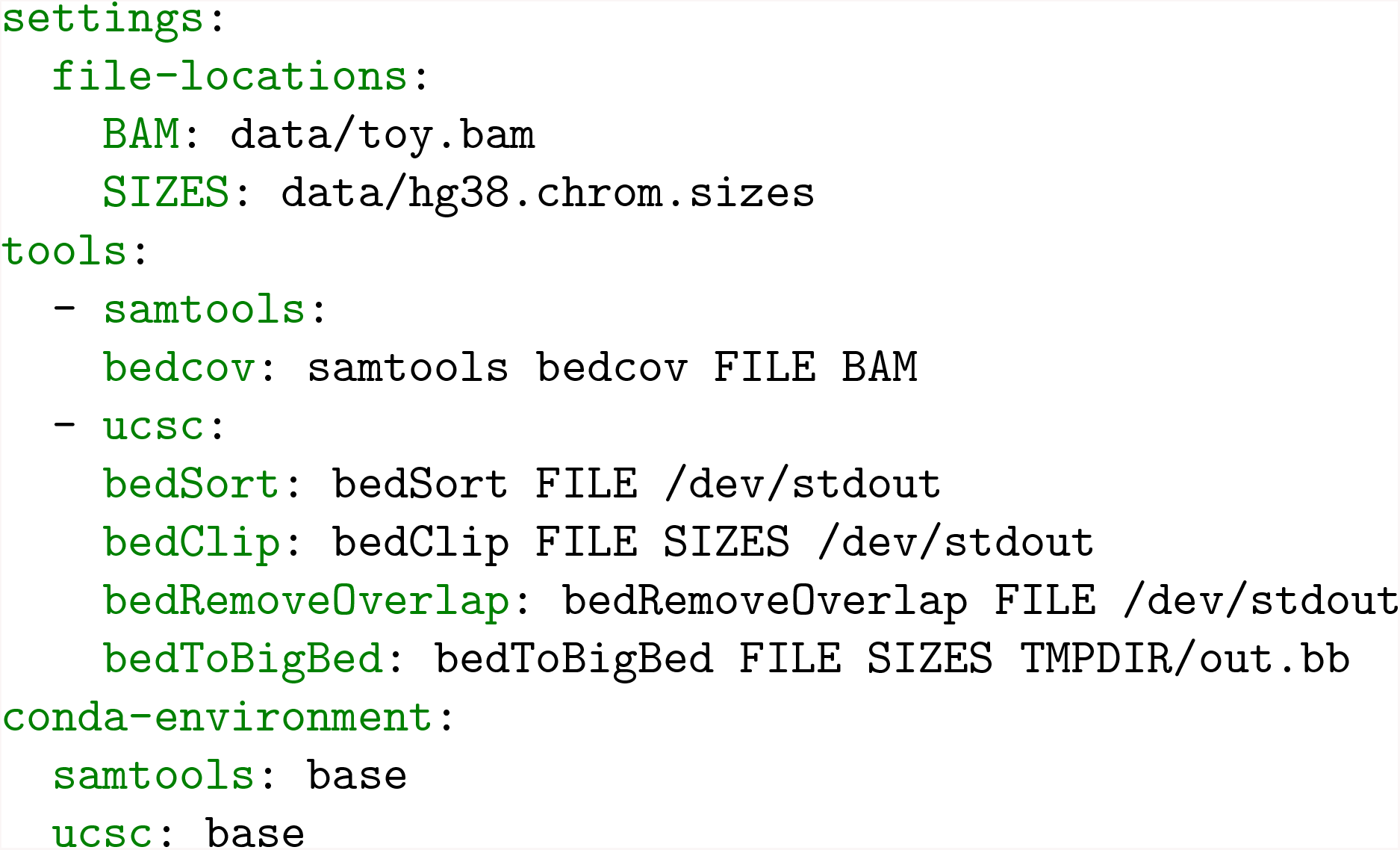
Excerpt from the Acidbio configuration file. The configuration file contains three sections. The “settings” section lists the location of files or directories that Acidbio will insert into command-line execution. The “tools” section contains the command-line invocations of the tested tools. In each invocation, Acidbio replaces “FILE” with the location of the test BED file. The “conda-environment” section lists each tool’s Conda environment name.

### 3.2 Tool discovery

To identify tools to test, we used Bioconda^10^, a repository that contains thousands of bioinformatics software packages. Each package contains one or more tools. We only included Bioconda packages with tools that have a command-line interface, as opposed to add-on modules executed within another program, and use the BED format as input. This excluded the numerous R, Bioconductor^12^, and Perl packages that have no command-line interface.

For packages that contain multiple tools, we selected a smaller set of subtools to test. We systematically identified these packages by manually examining the documentation for over 1000 packages to determine if it matches our criteria. We had to manually examine documentation because Bioconda has no structured metadata on each package’s input file formats. This process yielded 80 packages, with 99 tools total.

Some tools use the BED format as the primary input file, such as a mandatory argument. Examples include bedtools^23^ and high-throughput sequencing toolkits such as ngs-bits^98^. These tools generally perform calculations using the intervals found in the BED file.

Other tools use the BED format as a secondary input file, such as an optional argument. Tools that use BED as a secondary input file generally use it to define genomic intervals of interest for data in another file format, such as SAM. In the tools we tested, 60 packages used the BED format as the primary input file, and 20 packages used the BED format as a secondary input file.

After collecting a list of all the possible packages that we could test, we then attempted to install each package and run the tools. We excluded packages that we could not install or could not run without error on any input files. We found no cases where a package contained both working and broken tools.

### 3.3 Test suite

We created a test suite that contains tests for each BED*n* format, covering various edge cases drawn from our BED specification. The test suite contains both expected success test cases (Table 3) and expected fail test cases (Table 4). Some tests include validating ranges for numeric fields, validating character sets for alphanumeric fields, or data formatting for fields such as itemRgb or the block definitions.

**Table 3:**
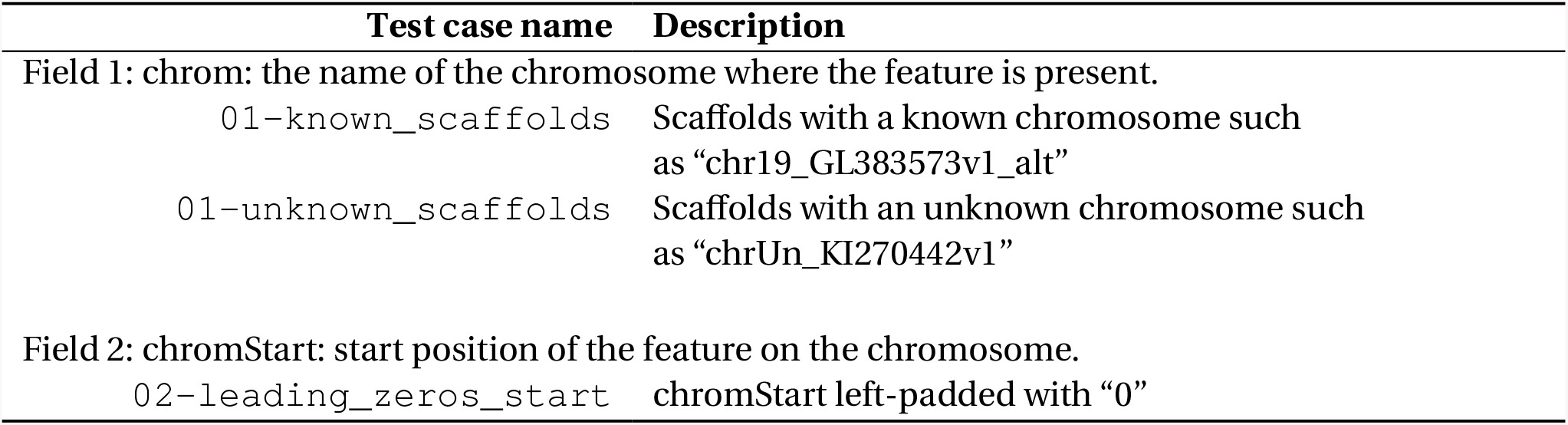
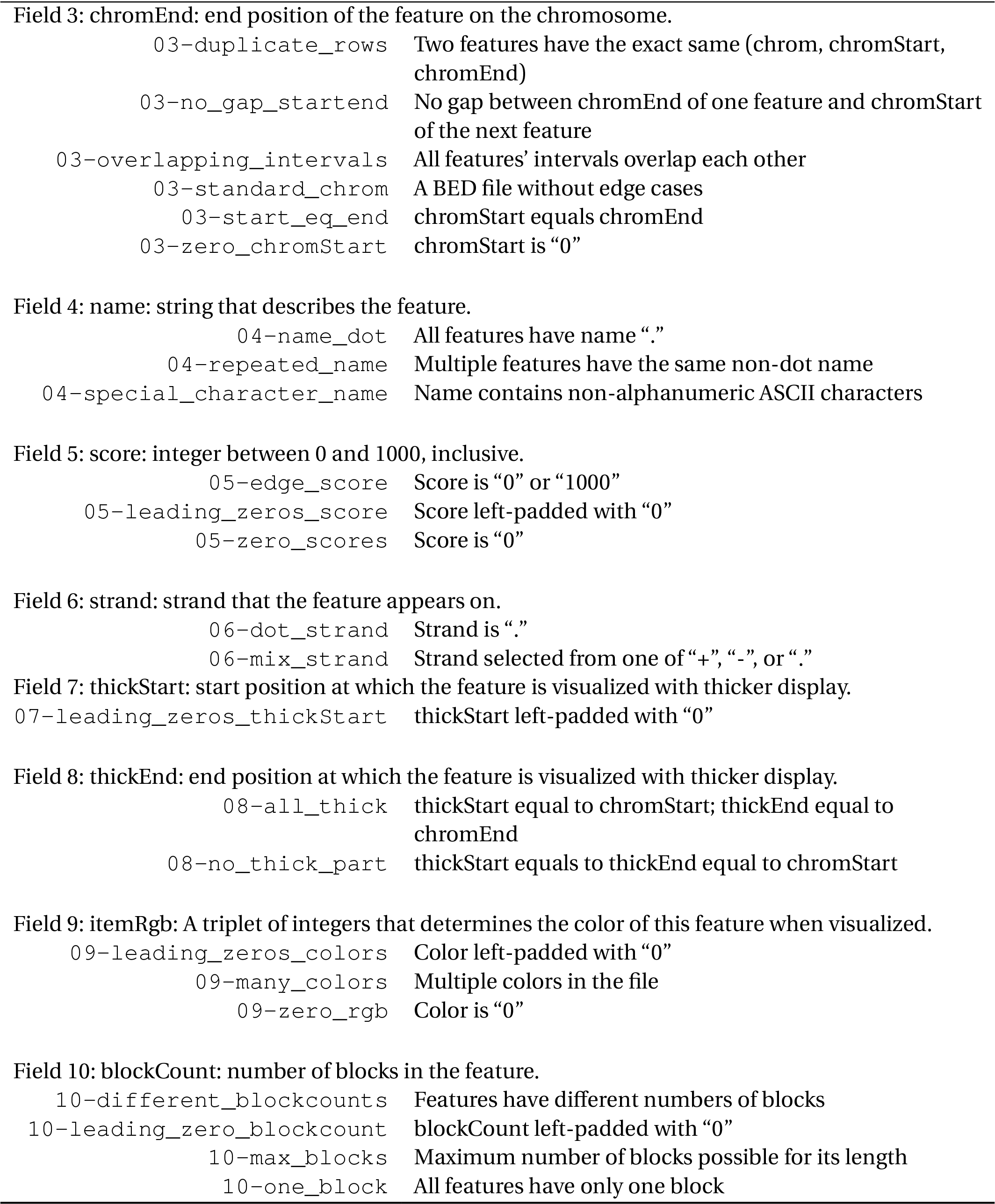
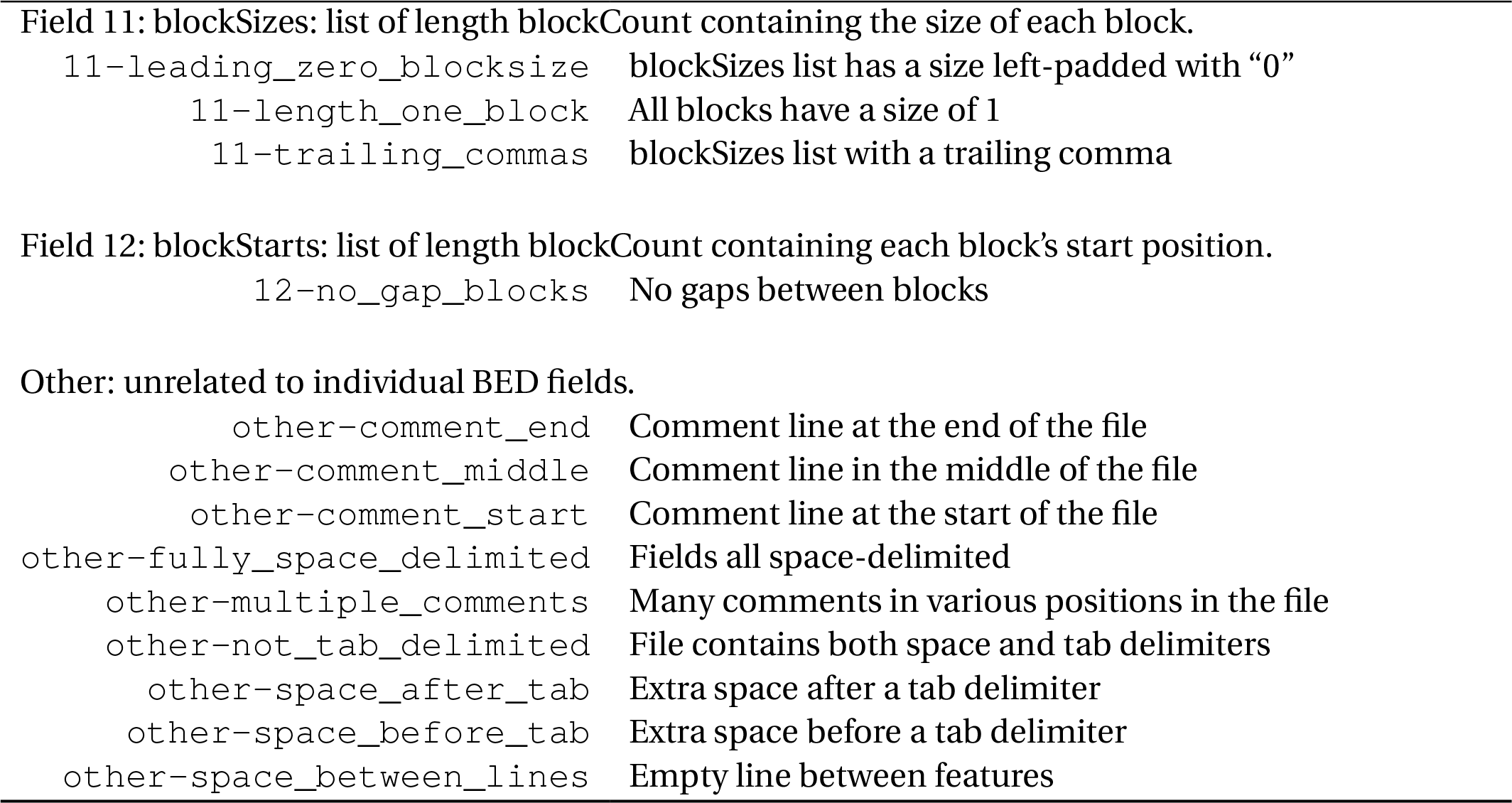
Expected success test cases.

| Test case name | Description |
| --- | --- |
| Field 1: chrom: the name of the chromosome where the feature is present. |  |
| 01-known_scaffolds | Scaffolds with a known chromosome such as “chr19_GL383573v1_alt” |
| 01-unknown_scaffolds | Scaffolds with an unknown chromosome such as “chrUn_KI270442v1” |
| Field 2: chromStart: start position of the feature on the chromosome. |  |
| 02-leading_zeros_start | chromStart left-padded with “0” |

**Table 3: Expected success test cases.** (Continued)
| Test case name | Description |
| --- | --- |
| Field 3: chromEnd: end position of the feature on the chromosome. |  |
| 03-duplicate_rows | Two features have the exact same (chrom, chromStart, chromEnd) |
| 03-no_gap_startend | No gap between chromEnd of one feature and chromStart of the next feature |
| 03-overlapping_intervals | All features' intervals overlap each other |
| 03-standard_chrom | A BED file without edge cases |
| 03-start_eq_end | chromStart equals chromEnd |
| 03-zero_chromStart | chromStart is "0" |
| Field 4: name: string that describes the feature. |  |
| 04-name_dot | All features have name "." |
| 04-repeated_name | Multiple features have the same non-dot name |
| 04-special_character_name | Name contains non-alphanumeric ASCII characters |
| Field 5: score: integer between 0 and 1000, inclusive. |  |
| 05-edge_score | Score is "0" or "1000" |
| 05-leading_zeros_score | Score left-padded with "0" |
| 05-zero_scores | Score is "0" |
| Field 6: strand: strand that the feature appears on. |  |
| 06-dot_strand | Strand is "." |
| 06-mix_strand | Strand selected from one of "+", "-", or "." |
| Field 7: thickStart: start position at which the feature is visualized with thicker display. |  |
| 07-leading_zeros_thickStart | thickStart left-padded with "0" |
| Field 8: thickEnd: end position at which the feature is visualized with thicker display. |  |
| 08-all_thick | thickStart equal to chromStart; thickEnd equal to chromEnd |
| 08-no_thick_part | thickStart equals to thickEnd equal to chromStart |
| Field 9: itemRgb: A triplet of integers that determines the color of this feature when visualized. |  |
| 09-leading_zeros_colors | Color left-padded with "0" |
| 09-many_colors | Multiple colors in the file |
| 09-zero_rgb | Color is "0" |
| Field 10: blockCount: number of blocks in the feature. |  |
| 10-different_blockcounts | Features have different numbers of blocks |
| 10-leading_zero_blockcount | blockCount left-padded with "0" |
| 10-max_blocks | Maximum number of blocks possible for its length |
| 10-one_block | All features have only one block |

**Table 3: Expected success test cases.** (Continued)
| Test case name | Description |
| --- | --- |
| Field 11: blockSize: list of length blockCount containing the size of each block. |  |
| 11-leading_zero_blocksize | blockSizes list has a size left-padded with "0" |
| 11-length_one_block | All blocks have a size of 1 |
| 11-trailing_commas | blockSizes list with a trailing comma |
| Field 12: blockStarts: list of length blockCount containing each block's start position. |  |
| 12-no_gap_blocks | No gaps between blocks |
| Other: unrelated to individual BED fields. |  |
| other-comment_end | Comment line at the end of the file |
| other-comment_middle | Comment line in the middle of the file |
| other-comment_start | Comment line at the start of the file |
| other-fully_space_delimited | Fields all space-delimited |
| other-multiple_comments | Many comments in various positions in the file |
| other-not_tab_delimited | File contains both space and tab delimiters |
| other-space_after_tab | Extra space after a tab delimiter |
| other-space_before_tab | Extra space before a tab delimiter |
| other-space_between_lines | Empty line between features |

**Table 4:** Expected fail test cases.

| Test case name | Description |
| --- | --- |
| Field 1: chrom: the name of the chromosome where the feature is present. |  |
| 01-missing_chrom | chrom is "." |
| 01-no_chrom | Missing chrom |
| 01-special_char_chrom | chrom has non-alphanumeric character |
| 01-quote_chrom | chrom enclosed with quotation marks |
| Field 2: chromStart: start position of the feature on the chromosome. |  |
| 02-float_start | chromStart is floating-point number |
| 02-no_start | Missing chromStart |
| 02-start_negative | chromStart is negative |
| 02-string_start | chromStart is not numeric |
| Field 3: chromEnd: end position of the feature on the chromosome. |  |
| 03-chromEnd_too_large | chromEnd exceeds maximum size of an unsigned integer |
| 03-end_negative | chromEnd is negative |
| 03-float_end | chromEnd is floating-point number |
| 03-no_end | Missing chromEnd |
| 03-start_gt_end | chromStart greater than chromEnd |
| 03-string_end | chromEnd is not numeric |

**Table 4: Expected fail test cases.** (Continued)
| Test case name | Description |
| --- | --- |
| Field 4: name: string that describes the feature. |  |
| 04-mixed_bed_types | Features have a different number of fields |
| 04-name_with_spaces | Name has a space |
| 04-no_name | Missing name |
| Field 5: score: integer between 0 and 1000, inclusive. |  |
| 05-float_score | Score is floating-point number |
| 05-no_score | Missing score |
| 05-score_gt_1000 | Score greater than 1000 |
| 05-score_lt_0 | Score less than 0 |
| 05-string_score | Score is not numeric |
| Field 6: strand: strand that the feature appears on. |  |
| 06-invalid_strand | Strand not one of "+", "-", or "." |
| 06-no_strand | Missing strand |
| Field 7: thickStart: start position at which the feature is visualized with thicker display. |  |
| 07-float_thickStart | thickStart is floating-point number |
| 07-no_thickStart | Missing thickStart |
| 07-string_thickStart | thickStart is not numeric |
| 07-thickStart_lt_chromStart | thickStart less than chromStart |
| Field 8: thickEnd: end position at which the feature is visualized with thicker display. |  |
| 08-float_thickEnd | thickEnd is floating-point number |
| 08-string_thickEnd | thickEnd is not numeric |
| 08-thickEnd_gt_chromEnd | thickEnd greater than chromEnd |
| 08-thickStart_gt_thickEnd | thickStart greater than thickEnd |
| 08-thickStartEnd_wide | Thick interval not contained within [chromStart, chromEnd) |
| Field 9: itemRgb: A triplet of integers that determines the color of this feature when visualized. |  |
| 09-incomplete_rgb | RGB value not a triplet or "0" |
| 09-no_rgb | Missing itemRgb |
| 09-string_rgb | itemRgb contains alphabetical characters |
| Field 10: blockCount: number of blocks in the feature. |  |
| 10-blockCount_negative | blockCount is negative |
| 10-blockCount_string | blockCount is non-numeric |
| 10-blockCount_zero | blockCount is "0" |
| 10-no_blockCount | Missing blockCount |

**Table 4: Expected fail test cases.** (Continued)
| Test case name | Description |
| --- | --- |
| Field 11: blockSizes: list of length blockCount containing the size of each block. |  |
| 11-block_outofrange | Block located outside the interval chromStart to chromEnd |
| 11-blocksizes_missing_comma | blockSizes list missing comma separators |
| 11-mismatch_count_long | blockSizes list longer than blockCount |
| 11-mismatch_count_short | blockSizes list shorter than blockCount |
| 11-size_negative | Block has negative size |
| Field 12: blockStarts: list of length blockCount containing each block's start position. |  |
| 12-bed13 | Feature has 13 fields |
| 12-overlapping_blocks | Blocks overlap each other |
| 12-start_outofrange | Block start position past chromEnd |
| 12-unsorted_block_starts | blockStarts list not sorted |
| Other: unrelated to individual BED fields. |  |
| other-quote_chrom | Feature enclosed in quotation marks |

We manually generated the test cases, designing them to make sense for all the tools tested. We used genomic intervals between positions 250000 and 260000 since one might find them in both chromosomes and non-chromosome scaffolds. Each test case varies based on the criteria tested. Some criteria only require a deviation in one field in one feature to generate a test case. For example, to test a score greater than 1000, only a single feature had a score greater than 1000. Other criteria required deviation in multiple features to generate a test case. For example, to test that the parser accepts strand “.”, we set all features to strand “.”.

We built tests upon each other—we repeated a test case for all BED variants with additional fields added. As an example, a test case in BED5 testing a negative score gets repeated in testing the BED6 through BED12 variants.

For tools that use BED as a secondary file format, we collected test files for their non-BED primary file formats. For each of these file formats, we sourced an example file from the creators of the format or from a repository such as a FASTA for GRCh38/hg38^109^ from the UCSC Genome Browser (https://hgdownload.cse.ucsc.edu/goldenpath/hg38/bigZips/). We edited non-BED files to ensure that their ranges matched the BED test cases. We also validated the collected non-BED files with a file validator, when possible.

Since the new formal BED specification prohibits BED10 and BED11, we considered all BED10 and BED11 tests expected fail, even if the test case fell under expected success for other BED variants.

### 3.4 Fuzzing

We used a fuzzing approach^110^ to automatically generate test cases beyond our manually designed test suite (Figure 7). We created an ANother Tool for Language Recognition 4 (ANTLR4) grammar^111^ to define the structure of the BED format and the possible values for each field. Then, we used a file generator that builds a file based on our grammar. We tested the tools using grammar-based fuzzing and grammarinator^112^ as the file generator.

**Figure 7:**
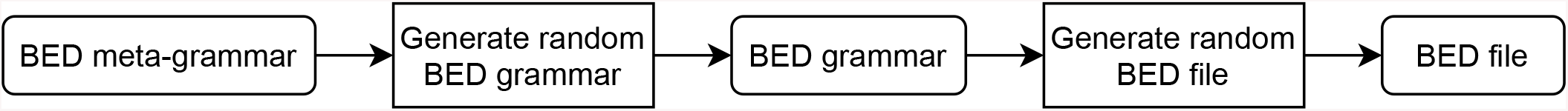
Flowchart depicting grammar-based fuzzing. Rounded rectangles: files; sharp rectangles: generators. The BED meta-grammar describes different BED grammars to introduce further variation in the generated BED file.

To introduce further variation into the BED file, we created an ANTLR4 meta-grammar that defines possible ANTLR4 BED grammars. The meta-grammar produces variation by allowing the BED grammar to vary on the structure or definition of fields. For example, the meta-grammar may produce a BED grammar that only allows tabs as the whitespace, or it may produce a BED grammar that allows both tabs and spaces. By varying the BED grammar produced, the user can test different combinations of field definitions and BED file structure that a single BED grammar cannot achieve.

### 3.5 Availability

Acidbio and the BED test suite are available at GitHub (https://github.com/hoffmangroup/acidbio) and deposited in Zenodo (https://doi.org/10.5281/zenodo.5784763). Results of each package on each test case and the scripts used to generate figures are available at Zenodo (https://doi.org/10.5281/zenodo.5784787).

## 4 Discussion

### 4.1 Use in software development

Acidbio can help researchers and programmers test their tools to improve the robustness and interoperability of their code. Acidbio can serve a similar function to the Web Standards Project Acid test suite^15^ designed to improve interoperability of web browsers. When the Web Standards Project created the Acid tests, many web browsers had poor compliance with existing web standards. Over time, browsers such as Opera^113^ and Internet Explorer^114^ began to achieve perfect performance on the Acid tests and interoperability improved. Similarly, we intend Acidbio to make it easier for developers to create bioinformatics software that more easily interoperates with other software.

To test new tools, developers need only create a short configuration YAML file to describe their tool’s command line interface, and run the Acidbio test harness. From the test results, a programmer may identify edge cases they missed and fix them before distributing their software. Once fixed, the programmer can put a BED badge in a software’s documentation to indicate that it interoperates with the BED format. Editors or reviewers of papers describing tools can use the test suite to verify the software’s quality. Package repository managers can also use the test suite to verify the quality of submitted packages.

### 4.2 The utility of a formal specification

The interpretation of a standard can turn into a matter of opinion. While formalizing the standard with a specification can help improve interoperability, the only way to truly ensure agreement on expected behavior involves further formalization through a formal grammar or including test cases in the standard. A deterministic grammar or test suite removes potential for misunderstandings about standard conformance.

### 4.3 Postel’s law

Postel’s law, “be conservative in what you do, be liberal in what you accept from others”^115^, related initially to how software sends and accepts messages over the internet. Adherence to Postel’s law helped the internet to succeed—leniency in accepting data without strict validation helped more organizations implement internet software^116^.

As seen here, many software tools have taken a liberal approach to accepting BED files. This seemingly increases the utility of these tools. By liberally accepting input, however, tools encourage BED producers to take a lackadaisical approach to correctness and interoperability, which leaves the format open to misuse^117^. Programmers may unwittingly create software that generates incorrect BED files if they only supply their output to downstream consumers with a liberal approach to validation. This results in technical debt, where problems lay undiscovered until after the developers complete the project, or years later, when it becomes much harder to fix.

The developers of the Extensible Markup Language (XML) format purposefully rejected Postel’s law, deciding that malformed XML files would raise fatal errors^118^. They did this because this approach encourages producers of the file format to strongly conform to the specification. A strict validation approach reduces opportunities for parsers to misunderstand input and prevents common errors from becoming accepted.

The lack of a strict validation approach for previous HyperText Markup Language (HTML) implementations led to a morass of incompatible and poorly described HTML file formats. This greatly increased the complexity of potential bugs in web browsers that could actually handle the existing base of web pages. Despite the existence of formal HTML specifications, web browsers had to create special “quirks modes” to handle HTML files that did not satisfy these specifications^119^.

The history of HTML and XML should inform file validation behavior in bioinformatics software. While one may not want to raise fatal errors for each non-conforming file, BED parsers must at least provide warnings when encountering them. Users can easily ignore warnings, however, or miss them in a stream of irrelevant and voluminous diagnostic information. To ensure that users notice problems with file formats and that programmers fix upstream generators, parsers must take a strict validation “warnings are errors” approach and refuse to parse invalid files.

### 4.4 Application to other bioinformatics file formats

Users and developers can apply the same methodology developed here to test other bioinformatics file formats for conformance. Establishing a common interface to parse a file format will improve interoperability of bioinformatics software and move closer to FAIR^6^ goals. For binary file formats or software written in languages with weak memory safety, testing and interoperability become even more important.

Computational tools described in scholarly papers often undergo precious little testing. The existence of test systems such as Acidbio make it easy to test that a tool interoperates with other software well. We recommend that when such a test system exists, journal editors, reviewers, and software repository managers ensure that the tool achieves good performance in the test suite prior to acceptance. After acceptance, managers can indicate which file formats the package uses as input and output to make searching for tools easier. Developers can also add badges similar to the BED badge to indicate software’s conformance to the relevant specification.

### 4.5 BED metadata

Tools parse BED files in the absence of in-band information embedded within the file. The lack of in-band information may lead to difficulties parsing BED files. For example, a tool cannot determine whether a BED file has custom fields without in-band information. With such metadata, tools can easily determine whether the input file has the fields it needs.

A header section at the beginning of a BED file can provide metadata to make parsing of BED files easier. The header can define the file’s BED variant and specify information such as the genome assembly used. For custom BED*n*+*m* files, the header can define the custom-defined fields, similar to the INFO lines in the VCF meta-information lines. Having a header would provide a direct method of supplying file metadata directly within the file, allowing parsers to easily read the BED file. Future versions of the GA4GH BED specification may add such metadata.

### 4.6 Limitations of the testing approach

Our testing approach applies the same BED files and secondary files to all the tools, except tools that use BAM input. Given the diversity of tools that use the BAM format, we could not find a single BAM file with data relevant to all tools. Instead, we used two different BAM files to avoid tools raising logical errors on our test cases.

Our testing approach only considers whether a BED parser accepts valid input and rejects invalid input. It does not consider correctness of the output. Developers can validate output file format using a file validation tool. For BED files, one can use bigToBigBed^18^ for file validation, keeping in mind the edge cases discussed above where its behavior differs from the GA4GH BED specification. Testing for correctness of analyses represents a much more difficult problem that one cannot trivially address.

The fuzzing approach also has some limitations. The quality of the generated test cases relies on the file generator to cover a wide range of possible BED files. For a grammar-based fuzzing approach, the grammar would have to describe all possible variations in a file, which becomes difficult for more complex file formats. Another potential issue with file generation arises if the generator has too few methods to vary its output files, generating files that do not cover enough cases. Machine learning or other approaches that inform future file generation from past unexpected behavior can address this issue^120^.

Other fuzzing approaches, such as mutation-based fuzzing, may not work in a bioinformatics context. Mutation-based fuzzers randomly modify existing files by adding random or nonsense characters. These fuzzers would not create diverse BED files and the mutations would likely create invalid and meaningless BED files. A security-oriented fuzzer such as American Fuzzy Lop^121^ can detect these vulnerabilities. Security-oriented fuzzers will produce test cases that can have nonsense data such as non-ASCII characters, which tests the tool’s ability to handle unexpected data.

## Acknowledgments

We thank Carl Virtanen (0000-0002-2174-846X) and Zhibin Lu (0000-0001-6281-1413) at the University Health Network High-Performance Computing Centre and Bioinformatics Core for technical assistance, Michael Hicks (University of Maryland, College Park; 0000-0002-2759-9223) and Leonidas Lampropoulos (University of Maryland, College Park; 0000-0003-0269-9815) for helpful discussions, and W. James Kent and the UCSC Genome Browser team for creating the BED format. This work was supported by the Natural Sciences and Engineering Research Council of Canada (RGPIN-2015-03948 to M.M.H.).

## Competing interests

The authors declare no competing interests.

## Author contributions

Conceptualization, D.D. and M.M.; Data curation, Y.N.; Formal analysis, Y.N.; Funding acquisition, M.M.H.; Investigation, Y.N., D.D., and E.G.R.; Methodology, Y.N. and M.M.H.; Project administration, M.M.H.; Resources, M.M.H.; Software, Y.N.; Supervision, D.D., E.G.R., and M.M.H.; Validation, Y.N.; Visualization, Y.N.; Writing — original draft, Y.N.; Writing — review & editing, Y.N., D.D., E.G.R., and M.M.H

## Notes

https://github.com/hoffmangroup/acidbio

https://doi.org/10.5281/zenodo.5784763

https://doi.org/10.5281/zenodo.5784787

### Competing Interest Statement

The authors have declared no competing interest.

## References

[1] Crouch et al. The Software Sustainability Institute: Changing research software attitudes and practices. Computing in Science Engineering, 15(6):74–80, November 2013.

[2] Mangul et al. Challenges and recommendations to improve the installability and archival stability of omics computational tools. PLOS Biology, 17(6):e3000333, June 2019.

[3] Schultheiss. Ten simple rules for providing a scientific web resource. PLOS Computational Biology, 7(5):e1001126, May 2011.

[4] Taschuk et al. Ten simple rules for making research software more robust. PLOS Computational Biology, 13(4):e1005412, April 2017.

[5] Karimzadeh et al. Top considerations for creating bioinformatics software documentation. Briefings in Bioinformatics, 19(4):693–699, July 2018.

[6] Wilkinson et al. The FAIR guiding principles for scientific data management and stewardship. Scientific Data, 3:160018, March 2016.

[7] Pauli. The basics of web hacking: tools and techniques to attack the web. Elsevier, 2013.

[8] Rehm et al. GA4GH: International policies and standards for data sharing across genomic research and healthcare. Cell Genomics, 1(2):100029, Nov 2021.

[9] Global Allicance for Genomics and Health. Genomic Data Toolkit. https://www.ga4gh.org/genomic-data-toolkit/.

[10] Grüning et al. Bioconda: sustainable and comprehensive software distribution for the life sciences. Nature Methods, 15(7):475–476, July 2018.

[11] Bioconda. Guidelines for Bioconda recipes. https://bioconda.github.io/contributor/guidelines.html.

[12] Gentleman et al. Bioconductor: open software development for computational biology and bioinformatics. Genome Biology, 5:R80, September 2004.

[13] Bioconductor. Bioconductor — package submission. https://www.bioconductor.org/developers/package-submission/.

[14] Knuth. A torture test for TeX. Technical report, Department of Computer Science, Stanford University, 1984.

[15] Hickson. Acid2. https://www.webstandards.org/files/acid2/test.html, 2005.

[16] Danecek et al. The variant call format and VCFtools. Bioinformatics, 27(15):2156–2158, August 2011.

[17] Yang et al. Scalability and validation of big data bioinformatics software. Computational and Structural Biotechnology Journal, 15:379–386, July 2017.

[18] Kent et al. BigWig and BigBed: enabling browsing of large distributed datasets. Bioinformatics, 26(17):2204–2207, July 2010.

[19] Haeussler et al. The UCSC genome browser database: 2019 update. Nucleic Acids Research, 47(D1):D853–D858, January 2019.

[20] Clawson. Personal communication, 2019.

[21] Li et al. The Sequence Alignment/Map format and SAMtools. Bioinformatics, 25(16):2078–2079, June 2009.

[22] Chen et al. Evolutionary analysis across mammals reveals distinct classes of long non-coding RNAs. Genome Biology, 17:19, February 2016.

[23] Quinlan et al. BEDTools: a flexible suite of utilities for comparing genomic features. Bioinformatics, 26(6):841–842, January 2010.

[24] Bioconvert. https://bioconvert.readthedocs.io/en/master/index.html, 2017.

[25] Khan et al. Intervene: a tool for intersection and visualization of multiple gene or genomic region sets. BMC Bioinformatics, 18:287, May 2017.

[26] Wang et al. RSeQC: quality control of RNA-seq experiments. Bioinformatics, 28(16):2184–2185, June 2012.

[27] Xu et al. A signal-noise model for significance analysis of ChIP-seq with negative control. Bioinformatics, 26(9):1199–1204, April 2010.

[28] Ramsköld et al. An abundance of ubiquitously expressed genes revealed by tissue transcriptome sequence data. PLOS Computational Biology, 5(12):e1000598, December 2009.

[29] Zerbino et al. WiggleTools: parallel processing of large collections of genome-wide datasets for visualization and statistical analysis. Bioinformatics, 30(7):1008–1009, December 2013.

[30] Mapleson et al. Efficient and accurate detection of splice junctions from RNA-seq with portcullis. GigaScience, 7(12):giy131, December 2018.

[31] Robinson et al. Integrative genomics viewer. Nature Biotechnology, 29(1):24–26, January 2011.

[32] Cooke et al. A unified haplotype-based method for accurate and comprehensive variant calling. Nature Biotechnology, 39(7):885–892, July 2021.

[33] Fang et al. An ensemble approach to accurately detect somatic mutations using SomaticSeq. Genome Biology, 16:197, September 2015.

[34] Shen et al. SeqKit: a cross-platform and ultrafast toolkit for FASTA/Q file manipulation. PLOS ONE, 11(10):e0163962, October 2016.

[35] Rausch et al. Alfred: interactive multi-sample BAM alignment statistics, feature counting and feature annotation for long- and short-read sequencing. Bioinformatics, 35(14):2489–2491, December 2018.

[36] Talevich et al. CNVkit: genome-wide copy number detection and visualization from targeted DNA sequencing. PLOS Computational Biology, 12(4):e1004873, April 2016.

[37] Mahony et al. An integrated model of multiple-condition ChIP-Seq data reveals predeterminants of Cdx2 binding. PLOS Computational Biology, 10(3):e1003501, March 2014.

[38] Heger et al. GAT: a simulation framework for testing the association of genomic intervals. Bioinformatics, 29(16):2046–2048, June 2013.

[39] Alneberg et al. Binning metagenomic contigs by coverage and composition. Nature Methods, 11(11):1144–1146, September 2014.

[40] Feng et al. PeakRanger: a cloud-enabled peak caller for ChIP-seq data. BMC Bioinformatics, 12:139, May 2011.

[41] Karunanithi et al. Automated analysis of small RNA datasets with RAPID. PeerJ, 7:e6710, April 2019.

[42] Heinz et al. Simple combinations of lineage-determining transcription factors prime cis-regulatory elements required for macrophage and B cell identities. Molecular Cell, 38(4):576–589, May 2010.

[43] Li et al. Measuring reproducibility of high-throughput experiments. Annals of Applied Statistics, 5(3):1752–1779, October 2011.

[44] Wang et al. CPAT: coding-potential assessment tool using an alignment-free logistic regression model. Nucleic Acids Research, 41(6):e74, January 2013.

[45] Langenberger et al. Evidence for human microRNA-offset RNAs in small RNA sequencing data. Bioinformatics, 25(18):2298–2301, July 2009.

[46] Hanghøj et al. DamMet: ancient methylome mapping accounting for errors, true variants, and post-mortem DNA damage. GigaScience, 8(4):giz025, April 2019.

[47] Herzeel et al. elPrep 4: A multithreaded framework for sequence analysis. PLOS ONE, 14(2):e0209523, February 2019.

[48] Heeringen et al. GimmeMotifs: a de novo motif prediction pipeline for ChIP-sequencing experiments. Bioinformatics, 27(2):270–271, November 2010.

[49] Zhang et al. Model-based analysis of ChIP-Seq (MACS). Genome Biology, 9:R137, September 2008.

[50] Thorvaldsdóttir et al. Integrative genomics viewer (IGV): high-performance genomics data visualization and exploration. Briefings in Bioinformatics, 14(2):178–192, April 2012.

[51] Feng et al. Regtools: Integrated analysis of genomic and transcriptomic data for discovery of mutations associated with aberrant splicing in cancer. Cancer Research, 78(13 Suppl):2285, July 2018.

[52] Kodali. cthreepo. https://github.com/vkkodali/cthreepo, 2020.

[53] Leonardi. Bedparse: feature extraction from BED files. The Journal of Open Source Software, 4(34):1228, February 2019.

[54] Boyle et al. F-seq: a feature density estimator for high-throughput sequence tags. Bioinformatics, 24(21):2537–2538, November 2008.

[55] Stovner et al. epic2 efficiently finds diffuse domains in ChIP-seq data. Bioinformatics, 35(21):4392–4393, March 2019.

[56] Cingolani et al. A program for annotating and predicting the effects of single nucleotide polymorphisms, SnpEff: SNPs in the genome of Drosophila melanogaster strain w1118; iso-2; iso-3. Fly, 6(2):80–92, April 2012.

[57] Picard toolkit. https://broadinstitute.github.io/picard/, 2019.

[58] Lopez et al. Explore, edit and leverage genomic annotations using Python GTF toolkit. Bioinformatics, 35(18):3487–3488, February 2019.

[59] Song et al. Identifying dispersed epigenomic domains from ChIP-Seq data. Bioinformatics, 27(6):870–871, March 2011.

[60] Gremme et al. GenomeTools: a comprehensive software library for efficient processing of structured genome annotations. IEEE/ACM Transactions on Computational Biology and Bioinformatics, 10(3):645–656, May 2013.

[61] Ay et al. Statistical confidence estimation for Hi-C data reveals regulatory chromatin contacts. Genome Research, 24(6):999–1011, June 2014.

[62] Daley et al. Modeling genome coverage in single-cell sequencing. Bioinformatics, 30(22):3159–3165, November 2014.

[63] Wala et al. VariantBam: filtering and profiling of next-generational sequencing data using regionspecific rules. Bioinformatics, 32(13):2029–2031, July 2016.

[64] Narzisi et al. Accurate de novo and transmitted indel detection in exome-capture data using microassembly. Nature Methods, 11(10):1033–1036, August 2014.

[65] Pongor et al. BAMscale: quantification of DNA sequencing peaks and generation of scaled coverage tracks. Epigenetics & Chromatin, 13:21, April 2020.

[66] Pedersen et al. Mosdepth: quick coverage calculation for genomes and exomes. Bioinformatics, 34(5):867–868, October 2017.

[67] Dale et al. Pybedtools: a flexible python library for manipulating genomic datasets and annotations. Bioinformatics, 27(24):3423–3424, September 2011.

[68] Willems et al. Genome-wide profiling of heritable and de novo STR variations. Nature Methods, 14(6):590–592, April 2017.

[69] Okonechnikov et al. Qualimap 2: advanced multi-sample quality control for high-throughput sequencing data. Bioinformatics, 32(2):292–294, January 2016.

[70] Neumann et al. Quantification of experimentally induced nucleotide conversions in high-throughput sequencing datasets. BMC Bioinformatics, 20:258, May 2019.

[71] Cingolani et al. Using Drosophila melanogaster as a model for genotoxic chemical mutational studies with a new program, SnpSift. Frontiers in Genetics, 3:35, March 2012.

[72] Costanza et al. A comparison of three programming languages for a full-fledged next-generation sequencing tool. BMC Bioinformatics, 20:301, June 2019.

[73] Breese et al. NGSUtils: a software suite for analyzing and manipulating next-generation sequencing datasets. Bioinformatics, 29(4):494–496, February 2013.

[74] Cretu Stancu et al. Mapping and phasing of structural variation in patient genomes using nanopore sequencing. Nature Communications, 8:1326, November 2017.

[75] Buske et al. Exploratory analysis of genomic segmentations with Segtools. BMC Bioinformatics, 12:415, October 2011.

[76] Sadedin et al. Bazam: a rapid method for read extraction and realignment of high-throughput sequencing data. Genome Biology, 20:78, 2019.

[77] Mikheenko et al. Versatile genome assembly evaluation with QUAST-LG. Bioinformatics, 34(13):i142–i150, June 2018.

[78] Neph et al. BEDOPS: high-performance genomic feature operations. Bioinformatics, 28(14):1919–1920, July 2012.

[79] Zhao et al. CrossMap: a versatile tool for coordinate conversion between genome assemblies. Bioinformatics, 30(7):1006–1007, December 2013.

[80] Pedersen et al. Comb-p: software for combining, analyzing, grouping and correcting spatially correlated p-values. Bioinformatics, 28(22):2986–2988, September 2012.

[81] Uren et al. Site identification in high-throughput RNA-protein interaction data. Bioinformatics, 28(23):3013–3020, December 2012.

[82] Sims et al. CGAT: computational genomics analysis toolkit. Bioinformatics, 30(9):1290–1291, January 2014.

[83] Kent et al. The human genome browser at UCSC. Genome Research, 12(6):996–1006, May 2002.

[84] Webster et al. Identifying, understanding, and correcting technical biases on the sex chromosomes in next-generation sequencing data. GigaScience, 8(7):giz074, July 2019.

[85] Ramírez et al. deepTools2: a next generation web server for deep-sequencing data analysis. Nucleic Acids Research, 44(W1):W160–W165, April 2016.

[86] Herzeel et al. elPrep: high-performance preparation of Sequence Alignment/Map files for variant calling. PLOS ONE, 10(7):e0132868, July 2015.

[87] Dunn et al. Plastid: nucleotide-resolution analysis of next-generation sequencing and genomics data. BMC Genomics, 17:958, November 2016.

[88] Farek. AlignStats. https://github.com/jfarek/alignstats, 2017.

[89] Hensly et al. atactk: a toolkit for ATAC-seq data. https://atactk.readthedocs.io/en/latest/index.html, 2015.

[90] Orchard et al. Quantification, dynamic visualization, and validation of bias in ATAC-Seq data with ataqv. Cell Systems, 10(3):298–306.e4, March 2020.

[91] Huddleston et al. Augur: a bioinformatics toolkit for phylogenetic analyses of human pathogens. Journal of Open Source Software, 6(57):2906, January 2021.

[92] Hof et al. Biopet: Towards scalable, maintainable, user-friendly, robust and flexible NGS data analysis pipelines. In 2017 17th IEEE/ACM International Symposium on Cluster, Cloud and Grid Computing (CCGRID), pages 823–829, 2017.

[93] Vorderman et al. chunked-scatter. https://github.com/biowdl/chunked-scatter, 2019.

[94] Heuer. dishevelled-bio. https://github.com/heuermh/dishevelled-bio.

[95] Kaul et al. Identifying statistically significant chromatin contacts from Hi-C data with FitHiC2. Nature Protocols, 15(3):991–1012, January 2020.

[96] Pertea et al. GFF utilities: GffRead and GffCompare. F1000Research, 9:304, April 2020.

[97] Curk et al. iCount: protein-RNA interaction iCLIP data analysis (in preparation), 2019.

[98] Sturm et al. ngs-bits short-read sequencing tools for diagnostics. In European Conference on Computational Biology, 2018.

[99] Kaul. Novasplice. https://aryakaul.github.io/novasplice/, 2018.

[100] Fang et al. Indel variant analysis of short-read sequencing data with Scalpel. Nature Protocols, 11(12):2529–2548, November 2016.

[101] Li. seqtk toolkit for processing sequences in FASTA/Q formats. https://github.com/lh3/seqtk, 2012.

[102] Pedersen. Smoove. https://github.com/brentp/smoove, 2018.

[103] Bentsen et al. ATAC-seq footprinting unravels kinetics of transcription factor binding during zygotic genome activation. Nature Communications, 11:4267, August 2020.

[104] Schiller. Data Biology: A quantitative exploration of gene regulation and underlying mechanisms. PhD thesis, University of California, San Francisco, 2013.

[105] Garrison. Vcflib: A C++ library for parsing and manipulating VCF files. https://github.com/ekg/vcflib, 2012.

[106] Bollen et al. sndrtj/wisestork: Version 0.1.0. https://doi.org/10.5281/zenodo.3245885, June 2019.

[107] McKeeman. Differential testing for software. Digital Technical Journal, 10(1):100–107, 1998.

[108] Godefroid et al. Grammar-based whitebox fuzzing. In Proceedings of the 29th ACM SIGPLAN Conference on Programming Language Design and Implementation, PLDI ‘08, pages 206–215, New York, NY, USA, 2008. Association for Computing Machinery.

[109] Schneider et al. Evaluation of GRCh38 and de novo haploid genome assemblies demonstrates the enduring quality of the reference assembly. Genome Research, 27(5):849–864, May 2017.

[110] Miller et al. An empirical study of the reliability of UNIX utilities. Communications of the ACM, 33(12):32–44, Dec 1990.

[111] Parr et al. Adaptive LL (*) parsing: the power of dynamic analysis. ACM SIGPLAN Notices, 49(10):579–598, October 2014.

[112] Hodován et al. Grammarinator: a grammar-based open source fuzzer. In Proceedings of the 9th ACM SIGSOFT International Workshop on Automating TEST Case Design, Selection, and Evaluation,pages 45–48, November 2018.

[113] Gohring. Acid test may prove new browsers are tough sell. https://www.networkworld.com/article/2309699/acid-test-may-prove-new-browsers-are-tough-sell.html, Mar 2006.

[114] Schofield. Internet Explorer 8 passes Acid2 test. https://www.theguardian.com/technology/blog/2007/dec/21/internetexplorer8passesaci?CMP=gu_com, Dec 2007.

[115] Postel et al. Transmission control protocol, Request For Comments 793. https://datatracker.ietf.org/doc/html/rfc793, 1981.

[116] Bray. On Postel, again. https://www.tbray.org/ongoing/When/200x/2004/01/11/PostelPilgrim, January 2004.

[117] Allman. The robustness principle reconsidered. Communications of the ACM, 54(8):40–45, August 2011.

[118] Bray. Dracon and Postel. https://www.tbray.org/ongoing/When/200x/2003/08/19/Draconianism, August 2003.

[119] Olsson. CSS properties. In CSS Quick Syntax Reference Guide, pages 43–45. Springer, 2014.

[120] Saavedra et al. A review of machine learning applications in fuzzing. arXiv, 1906:11133, June 2019.

[121] Zalewski. American Fuzzy Lop (2.52b). https://lcamtuf.coredump.cx/afl/, 2018.

